# Epigenetic Reprogramming of Tissue-Specific Transcription Promotes Metastasis

**DOI:** 10.1101/131102

**Authors:** Shuaishuai Teng, Yang Li, Ming Yang, Rui Qi, Yiming Huang, Qianyu Wang, Shasha Li, Kequan Lin, Yujing Cheng, Zhi John Lu, Dong Wang

## Abstract

Tumor metastasis is the cause of death for 90% of cancer patients, and no currently-available therapies target this multi-step process in which cancer cells spread from the local tissue of a primary tumor to distant organs where they establish secondary tumors^1^. Although epithelial-to-mesenchymal transition^2^, tumor-secreted exosomes^3^, epigenetic regulators as well as other genes^4-8^ have been implicated in metastasis, little is known about how cells adapt to and colonize new tissue environments. Here, we show that the epigenetics-mediated reprogramming of tissue-specific gene transcription in cancer cells promotes metastasis. Using colorectal cancer (CRC) as a model, we found in both clinical and cell line studies that metastatic CRC cells lose their colon-specific gene transcription program and gain a liver-specific gene transcription program as they metastasize in the liver. Further, we found this transcription reprogramming is driven by a reshaped epigenetic landscape of both typical and super-enhancers. Chemical inhibition of enhancer activity disrupts the ability of cells to execute altered transcription programs and consequently inhibits metastasis. Binding motif analysis of the enhancers in liver metastatic CRC cells identified the liver-specific transcription factor FOXA2 as a key regulator, and knocking down of *FOXA2* expression prevents the colonization of metastatic CRC cells in the liver of a mice xenograft model. These results, together with additional observations of similar reprogramming in several cohorts of clinical CRC tumor samples and in multiple other forms of metastatic cancers, indicate that this reprogramming may be a common feature of metastasis in multiple cancers and suggest the targeted disruption of this epigenetic reprogramming as a strategy for the development of therapies to treat metastasis, the leading cause of cancer-related mortality.

---

Tumor metastasis refers to the movement of tumor cells from a primary site to distant organs that they progressively colonize^9^. More than 100 years ago, Paget suggested the idea of metastasis as the interaction of “seeds” and “soil”^10^, but subsequent research has yielded only a limited understanding of the mechanism(s) through which metastatic cancer cells (“seeds”) adapt to and colonize a new tissue environment (“soil”), the crucial steps of the metastasis process^11^.

It has been reported that the expression of tissue-specific or cell-lineage genes, which are regulated by general and cell-type-specific transcription factors through regulatory genomic elements, such as enhancers, determines cell identity^12-15^ and may also promote macrophages adapting to a particular environment in innate immune system^16^.

To explore the importance of tissue-specific genes during metastasis, we initially evaluated the tissue-specific transcriptome profiles of primary and liver-metastatic CRC tumors in three publicly available CRC datasets. Compared with primary CRC tumors, liver metastatic CRC samples had up-regulated expression of some commonly-known liver-specific genes and down-regulated expression of commonly-known colon-specific genes (Fig. 1a, 1b). To further investigate these intriguing trends in gene transcription, we next performed an unbiased Gene Set Enrichment Analysis (GSEA)^17^ of these published datasets to identify the statistically significant differences in global gene expression between primary and liver-metastatic CRC tumors. We used the Genotype-Tissue Expression (GTEx, https://gtexportal.org/home/) classification to define tissue-specific gene signatures in this analysis. Consistent with our initial findings for commonly-known liver and colon genes, the GSEA showed that the set of significantly-upregulated genes in the liver-metastatic CRC tumors was significantly enriched for a liver-specific signature (FDR < 0.00001), and the set of genes with down-regulated transcription were significantly enriched for a colon-specific gene signature (FDR < 0.00001) (Fig. 1c, d, e). We confirmed that these enrichments were still significant even after removing up-regulated genes that could possibly have been introduced by contamination from normal liver tissues (Extended Data Fig. 1a, b, c). Strikingly, additional analyses of primary and metastatic tumors from other cancers using the same analysis and classification criteria revealed that this apparent transcriptional reprogramming also occurs in prostate-to-liver metastasis, prostate-to-bone metastasis, kidney-to-lung metastasis and breast-to-lymph-nodes metastasis (Extended Data Fig. 2a, b, c, d). These findings suggest that metastatic cells can lose the tissue-specific transcription program of the tissue from which they originate and gain the tissue-specific transcription program of a distant tissue. Moreover, this transcriptional reprogramming may be a common feature of metastasis in multiple types of cancer.

**Figure 1:**
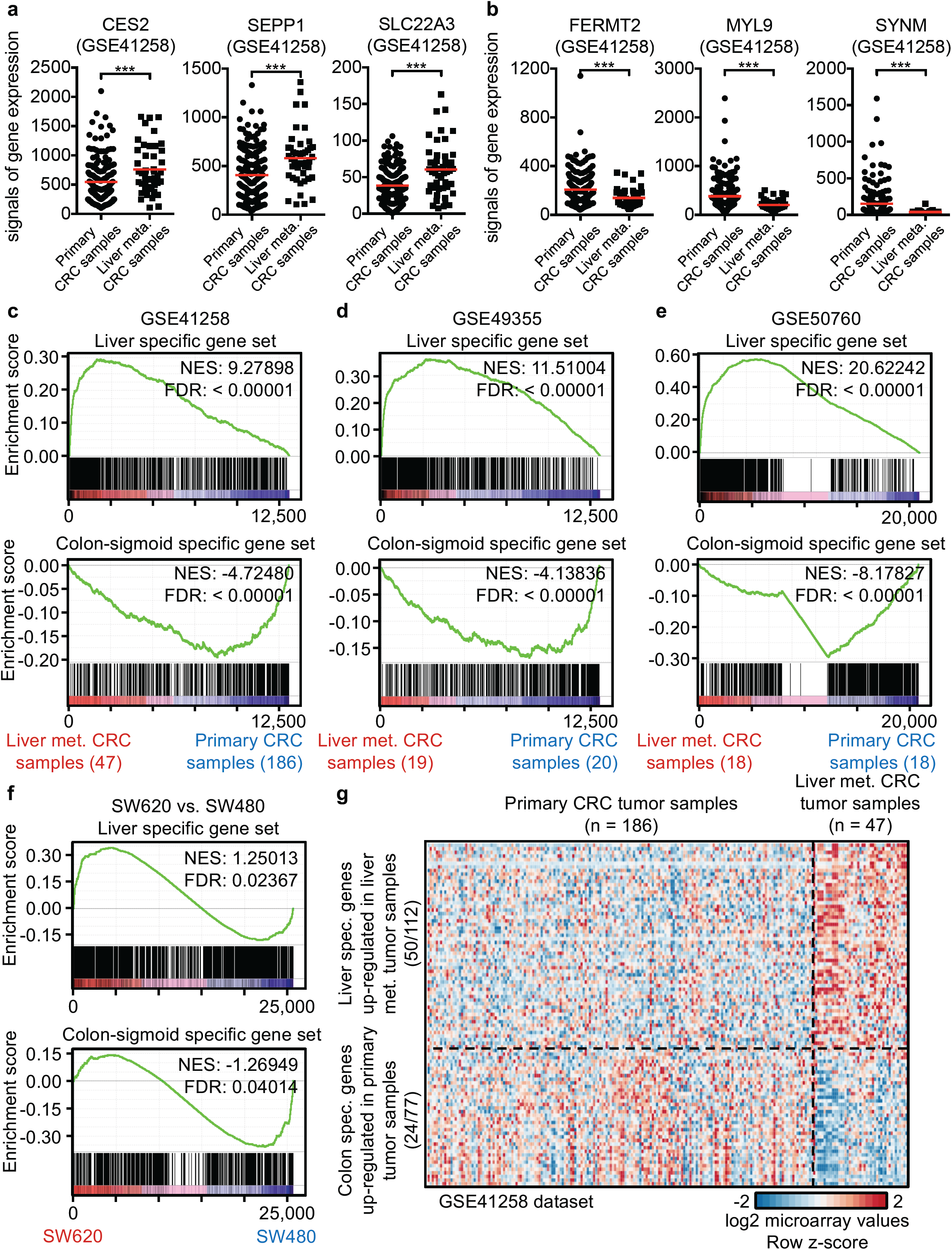
Reprogrammed tissue2specific transcription in liver metastatic CRC tumors and cell lines. **a,** Examples of liver-specific genes with significantly up-regulated transcription in liver metastatic CRC tumor samples. **b,** Examples of colon-specific genes with significantly down-regulated transcription in liver metastatic CRC tumor samples. The expression data are from the GSE41258 dataset. Each dot represents one primary or liver metastatic CRC tumorW red bars represent mean values. Statistically significant *P* values are indicated with asterisks (* *P* value < 0.01W ** *P* value < 0.005W *** *P* value < 0.001, by t-test). **c-e**, GSEA of liver-specific signatures (top) and colon-sigmoid-tissue specific signatures (bottom), as defined from the GTEx project database, in human primary and liver metastasis CRC tumors from the GSE41258 (**c**), GSE49355 (**d**), and GSE50760 (**e**) datasets. The genes are ranked by averaged expression values (log2 fold change) of multiple primary and liver metastasis CRC samples. The normalized enrichment scores (NES) and tests of statistical significance (FDR) are shown. **f**, GSEA of liver-specific signatures (top) and colonM sigmoid-tissue specific signatures (bottom) in SW620 and SW480 cells. The genes are ranked by the log2 fold change of the FPKM values in SW620 and SW480 cells. The NES and FDR are shown. **g**, Heatmap showing expression levels of liverM (50 out of 112 in **c**) and colon-specific genes (24 out of 77 in **c**) in human primary and liver metastatic CRC tumors.

Given the cellular heterogeneity of tumors, we sought to validate our initial findings in experiments using pure CRC cell lines (Extended Data Fig. 3a). SW620 cells were originally derived from a lymph node metastasis and can easily metastasize to liver in xenografts^18,19^. SW480 cells were derived from primary CRC tumor and show no ability to metastasize^18,19^. SW620 and SW480 cells were derived from the same patient^18^ thus share the same genetic background, providing an appropriate model to study reprogrammed transcription. RNA-seq gene expression profiling revealed striking differences between non-metastatic SW480 and liver-metastatic SW620 cells (Extended Data Fig. 3b). GSEA of our RNA-seq data for the two cell lines showed that a liver-specific gene signature was enriched in the significantly up-regulated genes in SW620 cells (FDR = 0.02), while a colon-specific gene signature was enriched in the significantly down-regulated genes (FDR = 0.04) (Fig. 1f). Specifically, SW620 cells had up-regulated expression of 112 liver-specific genes and down-regulated expression of 77 colon-specific genes (Extended Data Fig. 3c, d). Encouragingly, about half of the up-regulated liver-specific genes and about one third of the down-regulated colon-specific genes from the analysis of these two cell lines had similar expression trends in primary and liver-metastatic CRC tumor samples (Fig. 1g). Collectively, our results demonstrate that both liver metastatic CRC tumors and CRC cell lines gain a liver-specific transcription program and lose a colon-specific transcription program as these cells undergo metastasis.

Enhancers have been implicated in the regulation of tissue-specific or cell-lineage gene expression^12^. We therefore used chromatin immunoprecipitation sequencing (ChIP-seq) to investigate variations in the enhancer landscapes between SW620 and SW480 cells. Analysis using two antibodies against enhancer-specific histone modifications (H3K27ac and H3K4me2) showed that the deposition patterns of H3K27ac and H3K4me2 differed significantly between SW620 and SW480 cells (Fig. 2a and Extended Data Fig. 4a, b). An integrated analysis of our RNA-seq and ChIP-seq datasets showed that genes with up-regulated mRNA expression tended to be enriched in H3K27ac and H3K4me2 deposition near their genomic loci in SW620 cells (Fig. 2b and Extended Data Fig. 4c). Analysis of liver-specific gene expression in SW620 cells also showed that the genome regions around loci encoding up-regulated liver-specific genes had enriched H3K27ac (*P* value = 2.2e-16, by t-test) and H3K4me2 (*P* value = 2.2e-16, by t-test) deposition (Fig. 2c).

**Figure 2:**
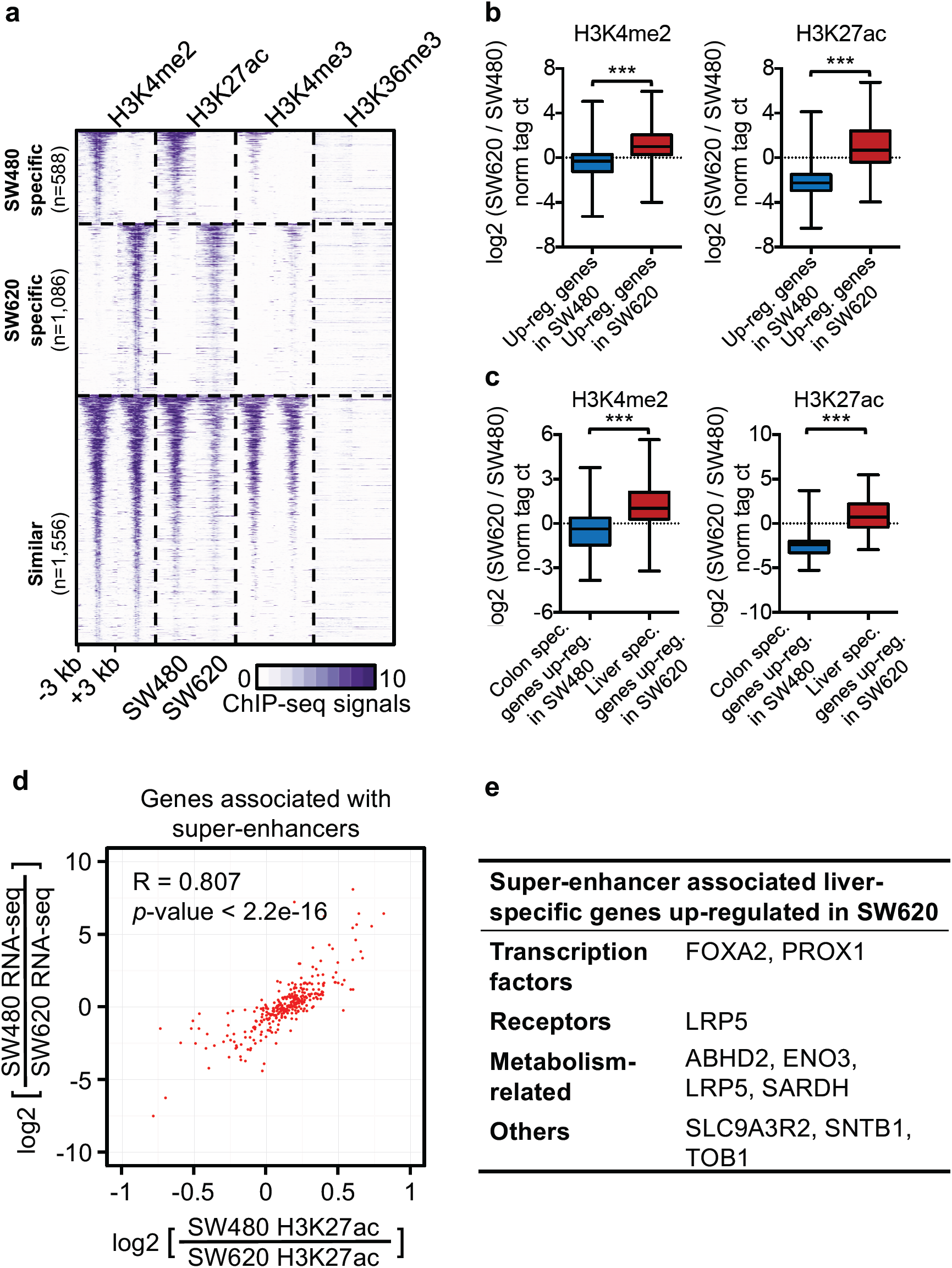
Variation in enhancer the landscape between primary and liver2metastatic CRC cell lines. **a,** Distribution of histone marks within ± 3 kb windows around the genomic regions in SW620 cells (n=1,086) and SW480 cells (n=588). The ChIPMseq datasets for H3K4me2, H3K27ac, H3K4me3, and H3K36me3 were each aligned with respect to the center of the H3K4me2 signal, and were sorted by the length of the H3K4me2Mmarked regions. ChIPMseq signals are tag counts normalized to 1×10^7^ uniquely mapped tags for each bin. **b**, Boxplots showing log2 ratios of SW620 to SW480 tag densities for genomic regions marked by H3K4me2 and H3K27ac around genes that are highly expressed in SW480 and SW620 cells, respectively. Statistically significant *P* values are indicated with asterisks (*** *P* value < 0.001, by Welch’s t-test). **c**, Boxplots showing log2 ratios of SW620 to SW480 tag densities for genomic regions marked by H3K4me2 and H3K27ac around un-regulated colon-specific genes in SW480 cells and up-regulated liver-specific genes in SW620 cells. Statistically significant *P* values are indicated with asterisks (*** *P* value < 0.005, by t-test). **d**, Scatterplot of the relationship between the ratio of SW480 to SW620 H3K27ac tag density at superM enhancers (x axis) and the ratio of nearest gene expression (y axis). The RNAMseq and ChIPMseq signals are the log2 of tag counts normalized to 1×10^7^ uniquely-mapped tags. The Pearson correlation coefficient is 0.807 and *P* < 2.2e-16. **e**, Examples of liver-specific genes associated with super-enhancers in SW620 cells.

So-called super-enhancers are a small fraction of total enhancers and encompass broad chromatin domains with H3K27ac deposition near genes essential for defining cell identity^20,21^. By identifying super-enhancers in SW480 and SW620 cells based on our H3K27ac ChIP-seq datasets, we found that in addition to the 264 super-enhancers common to both cell lines, there are 280 and 215 unique super-enhancers in SW620 and SW480 cells, respectively (Extended Data Fig. 4d). Comparison between our ChIP-seq and RNA-seq data revealed a high Pearson correlation coefficient (R = 0.807, *P* value < 2.2e-16) between the genome-wide distribution of super-enhancers and the expression levels of the nearest genes in these two cell lines (Fig. 2d and Extended Data Fig. 4e). Notably, many of the liver-specific genes in SW620 cells were found near SW620-unique super-enhancers; while the colon-specific genes in SW480 cells were found close to SW480-unique super-enhancers (Fig. 2e and Extended Data Fig. 4f, g). Our epigenetic and transcriptomics experiments demonstrate that reshaped enhancer and super-enhancer landscapes are involved in the reprogramming of tissue-specific transcription in CRC cells during metastasis.

A small molecule bromodomain inhibitor JQ1 can disrupt the binding of bromodomain containing 4 (BRD4) to enhancers^22^ and thus inhibits enhancer activity. We used JQ1 to disrupt the influence of enhancers on gene transcription in SW620 cells to explore the role of reshaped enhancer landscapes in the reprogramming of tissue-specific transcription in metastatic CRC cells. JQ1 treatment of SW620 cells resulted in the down-regulation of a set of liver-specific genes (Fig. 3a, b). GSEA further confirmed the liver-specific gene signature was significantly enriched in the set of down-regulated genes from JQ1-treated SW620 cells (FDR = 0.005) (Fig. 3c). Further, we noted the SW620-unique enhancers are more enriched near the genes that were down-regulated by JO1 treatment than common enhancers (*P* value = 4.148e-06, Fisher’s exact test) (Fig. 3d). Our JQ1 treatment results thus indicate that reprogramming of transcription in metastatic CRC cells is driven by reshaped enhancer landscapes. BRD4 inhibitors have been reported to suppress CRC cells metastasizing to the liver in mice, but very little is known about any related mechanisms of action^23^. Our findings suggest that the anti-metastasis effect of BRD4 inhibition may be conferred by preventing metastatic CRC cells from executing the liver-specific transcription program that is specified and driven by an altered enhancer landscape.

**Figure 3:**
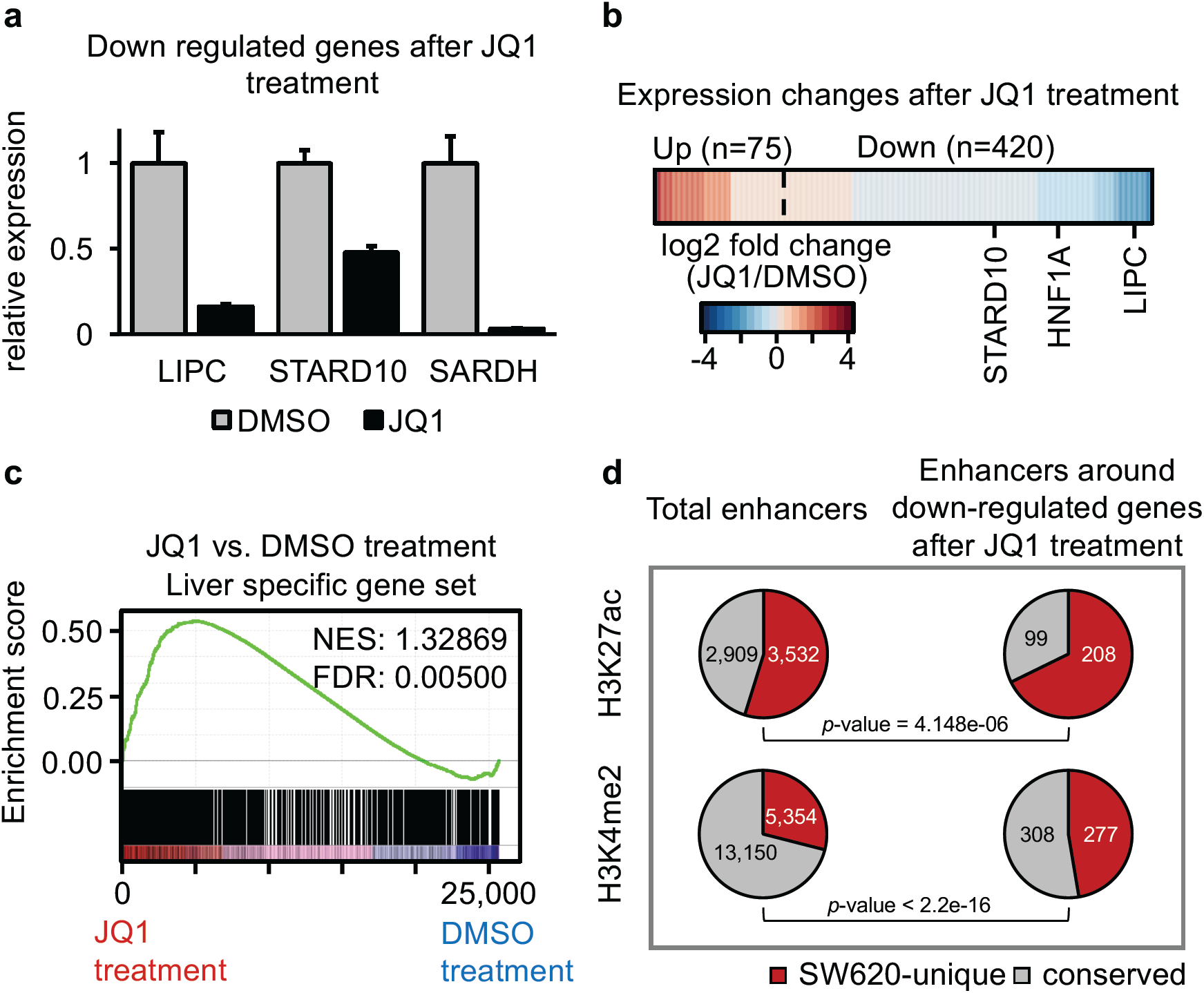
Chemical disruption of enhancer activity inhibits the liver2specific gene expression program. **a**, JQ1 treatment reduces the expression of a subset of liver-specific genes in SW620 cells. qRTMPCR results are presented for the expression of three genes. Error bars represent the standard deviation (SD) of three biological replicates. **b**, Heatmap showing RNAMseq analysis of JQ1Mtreated SW620 cellsW examples of significantly down-regulated liver-specific genes are labeled. The log2 fold change of normalized tag counts between JQ1Mtreated and DMSOMtreated (control) cells are plotted for differentially expressed genes (2 fold or greater changeW FDR ≤ 0.05). **c**, GSEA analysis of the liver-specific gene set (GTEx) in control and JQ1M treated SW620 cells. NES and FDR are shown. **d**, The expression of genes near SW620Munique enhancers are significantly more sensitive to JQ1 treatment than genes with conserved enhancers marked by H3K27ac or H3K4me2. The same trend was also found in super-enhancers marked by H3K27ac. The *P* values are from Fisherșs exact test.

We conducted a *de novo* binding motif analysis that examined all of the unique enhancers in SW620 cells (4,635) and found that the FOXA binding motif was the most highly-enriched (Fig. 4a and Extended Data Fig. 5a). There are three members of the *FOXA* gene family, and *FOXA2* is a well-known liver lineage-determining transcription factor^24^ that was highly expressed in SW620 cells (Extended Data Fig. 5b). ChIP-seq analysis with an antibody against FOXA2 showed that in SW620 cells FOXA2 occupied 35% of the SW620-unique enhancers but only occupied 0.01% of the SW480-unique enhancers (Fig. 4b and Extended Data Fig. 5c). Moreover, the deposition of H3K4me2 around FOXA2 binding sites was also significantly stronger in SW620 cells than in SW480 cells (Extended Data Fig. 5d). shRNA-based knocking-down of *FOXA2* expression in SW620 cells (Extended Data Fig. 6a) resulted in significant down-regulation of 624 genes (> 50% decrease; FDR ≤ 0.05) (Fig. 4c), and these genes were generally located near FOXA2-bound enhancers (*P* value = 2.2e-16, by t-test) (Extended Data Fig. 6b). Considering the primary functions of the liver, we were encouraged to find that a gene ontology analysis showed the top-ranked enriched term among the down-regulated genes upon FOXA2 knocking down was ‘metabolic process’ (Extended Data Fig. 6c). These results establish that FOXA2 binds to the SW620-unique enhancers in liver-metastatic CRC cells and functionally activates a set of liver-specific genes.

**Figure 4:**
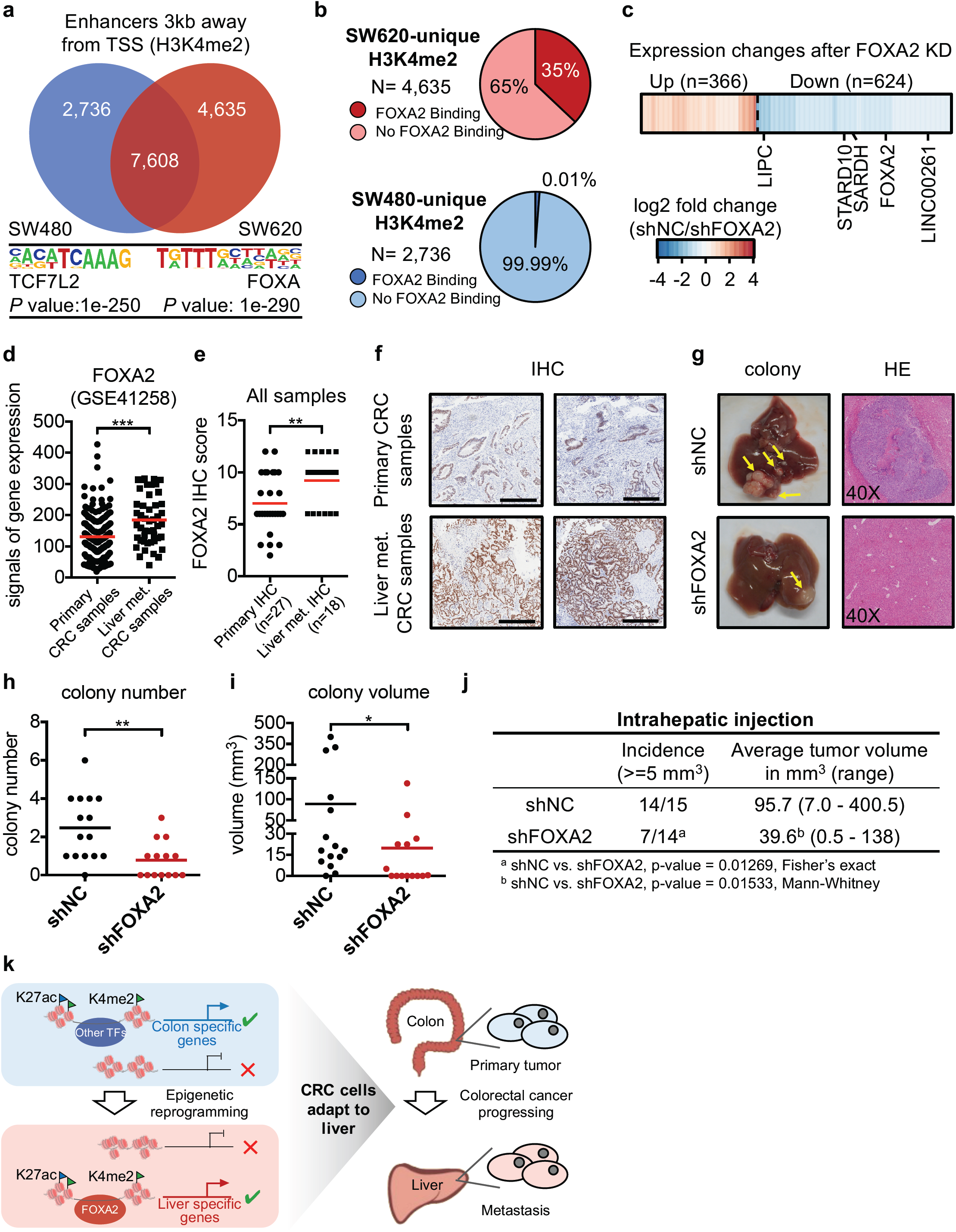
FOXA2 binds to the SW6202unique enhancers and promotes CRC liver metastasis. **a**, Enriched DNA motifs with significant *P* values, and the top DNA binding motif identified in a *de novo* motif analysis of non-promoter regions marked by H3K4me2 (far from 3 kb upM or downstream of TSS) in SW480 and SW620 cells. **b**, Percentage of FOXA2 binding sites within SW620Munique H3K4me2Mmarked enhancer regions (1636 out of 4,635) (top) and SW480Munique H3K4me2Mmarked enhancer regions (38 out of 2,736) (bottom) in SW620 cells. **c**, Heatmap showing RNAMseq analysis of SW620 cells with *FOXA2* knockdown. The log2 fold change of normalized tag counts between JQ1M and DMSOMtreated cells are plotted for differentially expressed genes (> 50% decrease or increaseW FDR ≤ 0.05). Examples of significantly downM regulated liver-specific genes in FOXA2 knockdown cells. **d**, *FOXA2* is highly expressed in human liver metastatic CRC tumors as compared with primary tumors. The expression data are from the GSE41258 dataset. Each dot represents one primary or liver metastatic CRC tumor, and the red bar represents the mean value. Statistically-significant *P* values are indicated with asterisks (*** *P* value < 0.001, by t-test). **e**, Plots showing the IHC scores for the nuclear FOXA2 in human primary CRC tumors and liver metastases. Each dot represents one primary or liver metastatic CRC tumor, and the red bar represents the mean value. Statistically-significant *P* values are indicated with asterisks (** *P* value < 0.005, by t-test). **f**, Representative IHC images of FOXA2 expression in paired CRC primary tumors and liver metastases. Scale bars, 500 μm. **g**, Representative light images (left) and HE images (right) of liver metastases 6 weeks after intrahepatic injection in mice. **h, i**, Both the number (b) and the volume (c) of the liver metastases were significantly reduced in response to FOXA2 knockdown after intrahepatic injection. Each dot represents a mouse (* *P* value < 0.05, ** *P* value < 0.005, by t-test). **j**, Incidence and average tumor volume of liver metastases 6 weeks after intrahepatic injection. **k**, Model for epigenetic reprogramming of tissue-specific transcription that promotes CRC metastasis to the liver.

We analyzed published gene expression datasets and found that *FOXA2* was more highly expressed in human CRC liver metastases than in primary CRC tumors (Fig. 4d). To exclude the possibility that contamination from normal liver tissue may be responsible for the increased *FOXA2* expression in the tumor samples, we assessed FOXA2 protein levels by immunohistochemistry (IHC) in a tissue microarray containing 18 liver metastases and 27 primary CRC tumors. This result clearly showed that the FOXA2 protein levels are significantly higher in liver metastatic CRC cells than in primary CRC cells (*P* value = 0.004, by t-test) (Fig. 4e, f and Supplementary Table 3). When we analyzed published clinical data, we found that the subset of CRC patients with high expression levels for FOXA2-targeted genes (91 genes, Supplementary Table 4) in their primary tumors had significantly worse overall survival (*P* value = 0.006, by log-rank test) and recurrence free survival (*P* value = 0.007, by log-rank test) outcomes than did a subset of patients with low expression levels for these genes (Extended Data Fig. 6d), strongly supporting an important role for FOXA2 in CRC metastasis. Strikingly, our data revealed that the liver-specific gene *LIPC* was directly up-regulated by FOXA2 (Extended Data Fig. 6e, f), which is consistent with its high expression in CRC liver metastases and the established function in promoting CRC tumor metastasis to liver^25^.

To investigate whether knocking-down *FOXA2* expression can alter the adaptation and colonization of metastatic CRC cells in the liver, we used a xenograft model to examine the metastatic potential^26^ of *FOXA2*-shRNA SW620 cells and control-shRNA SW620 cells in mice. Six weeks after intrahepatic-injection-based implantation of cells in the livers of nude mice, we found that tumors expressing control shRNA had a large number and big (macroscopically visible) metastases in liver whereas tumors expressing *FOXA2* shRNA had significantly fewer metastases that were also much smaller (Fig. 4g-j). Importantly, knocking-down of *FOXA2* in SW620 cells did not inhibit cell proliferation in cultures (Extended Data Fig. 7a) and did not inhibit the growth of subcutaneous tumors in nude mice (Extended Data Fig. 7b, c), indicating that the promotion function of FOXA2 on CRC tumor growth is context-dependent, e.g. only occurs in the liver. Our results clearly demonstrate that knocking-down of FOXA2 inhibits liver-specific gene transcription, thereby suppressing the adaptation and colonization of metastatic CRC cells in the liver.

We propose an ‘original to distant transition’ (ODT) working model to explain our observations about the epigenetic reprogramming of tissue-specific transcription that occurs in liver metastatic CRC cells (Fig. 4k). This model can explain how the CRC cells from the colon can adapt to and colonize distant liver tissue. Importantly, the ODT model predicts that there are multiple stages of the reprogramming process that could be targeted therapeutically, including the enzymes responsible for reshaping epigenetic landscapes and/or the transcription factors or other regulators that function in the altered transcription programs.

This ODT process appears to occur in multiple cancer types, and similar therapeutic strategies should in theory also work in other cancers. Recalling that there are at present no therapies to treat metastasis, and considering that this process causes the death of 90% of cancer patients, the mechanistic insights from our study suggest a timely new conceptual framework for scientists and clinicians to use as they seek to detect, treat, and perhaps even prevent metastasis.

## Acknowledgements

This work was supported by a grant from the National Natural Science Foundation of China (81673460) and fundings from Tsinghua-Peking Joint Center for Life Sciences and Beijing Municipal Science & Technology Commission. We thank Xuerui Yang (Tsinghua University, China) for help with cell proliferation assay. We thank Qian Gao for help with HiSeq 1500 operation. We thank members of the Tsinghua University Animal Resources Center for their care of the mice. The results published here are in whole or part based upon data generated by The Cancer Genome Atlas managed by the NCI and NHGRI. Information about TCGA can be found at http://cancergenome.nih.gov.

## Author contributions

D.W. and Z.J.L. conceived and supervised the project. S.S.T. performed the experiments with the help of M.Y., Q.Y.W. and Y.J.C.. Y.L. performed the computational analyses with the help from R.Q., Y.M.H. S.S.L. and Q.K.L.. D.W. and Z.J.L. wrote the manuscript with contributions from all other authors.

## METHODS

### Cell culture

SW480 and SW620 cell lines were obtained from China Infrastruture of Cell Line Resources and cultured as described^27^. Briefly, SW480 and SW620 cells were cultured in DMEM (Gibco Cat. No. C11995500BT) supplemented with 10% fetal bovine serum (Gemini Cat. No. 900-108) and 1% penicillin/streptomycin (Gibco Cat. No. 15140-122). Cells were cultured at 37°C with 5% CO_2_. Cells used to inject mice were stably transfected with luciferase.

### Plasmids and shRNA infections

A lentiviral U6-based expression vector containing PuroR-T2A-mCherry was used to express shRNAs. The lentiviral vector was digested by BsmbI, followed by annealed shRNA oligos insertion, to clone shRNA expression plasmids. We used two shRNA targeting sites against FOXA2, as follows: FOXA2-shRNA1: 5’ GAACGGCATGAACACGTACAT 3’ (from Sigma-Aldrich Corporation, TRCN0000014915); FOXA2-shRNA2: 5’ GCAAGGGAGAAGAAATCCATA 3’, as previously described^28^. shControl: 5’ CAACAAGATGAAGAGCACCAA 3’ Lentiviral vector particles were produced by tri-transfection of plasmids harboring the packaging construct, the transfer vector and the envelope-expressing construct into 293T cells using DNAfect reagents (Cwbio Cat. No. CW0806). Viral supernatants were harvested and used for infections or stored at -80°C. Stable FOXA2 knockdown cell lines were generated by using lentiviral U6-based expression vectors. Stable populations were selected with 2 μg/ml puromycin (Sigma-Aldrich Cat. No. P9620). Knockdown was confirmed by RT-qPCR and western blotting.

### Western Blotting

Cells were lysed in RIPA buffer. Proteins were separated by 10% polyacrylamide gel and transferred to polyvinylidene membranes (Bio-Rad Cat. No. 170-4159), which were blocked for one hour at room temperature in TBS with Tween 20 (TBST) containing 5% BSA and subsequently probed with primary antibodies overnight at 4°C. After incubating the membrane with anti-rabbit peroxidase-conjugated secondary antibody (Cell Signaling Technology Cat. No. 7074), protein levels were detected with SuperSignal West Pico reagents (ThermoFisher Scientific Cat. No. 34095). Primary antibodies were prepared in 5% BSA in TBST. The following primary antibodies were used:FOXA2 (Abcam Cat. No. ab108422), GAPDH (Cell Signaling Technology Cat. No. 2118)

### SYBR-Green Real-Time PCR

Total RNA was extracted from cells by using TRIzol® Reagent (Ambion Cat. No. 15596018), and reverse-transcribed to cDNA by using the Thermo Scientific RevertAid First Strand cDNA Synthesis Kit (ThermoFisher Scientific Cat. No. K1622). Quantitative PCR (qPCR) involved use of the KAPA SYBR FAST Universal qPCR Kit (KAPA Cat. No. KK4601) in triplicate, with normalization to GAPDH. Primer sequences (from 5’ to 3’) are listed in follow: FOXA2 forward primer, TTGCTGGTCGTTTGTTGTGG; FOXA2 reverse primer, GTTCATGTTGCTCACGGAGG; LIPC forward primer, GTTCATGTTGCTCACGGAGG; LIPC reverse primer, GGCTGAAGCTGTTCATGTCA; STARD10 forward primer, GGCTGAAGCTGTTCATGTCA; STARD10 reverse primer, TTCCACTCGGGGTACTTGAG; SARDH forward primer, TTCCACTCGGGGTACTTGAG; SARDH reverse primer, AGCCCCACCAGGTAGAACTT; GAPDH forward primer, AGCCCCACCAGGTAGAACTT and GAPDH reverse primer, AGCCTTCTCCATGGTGGTGAAGAC.

### Proliferation assay

The effect of FOXA2 knockdown on SW620 cell proliferation was monitored in real-time by using the Incucyte Live-Cell Imaging System (Essen BioScience, USA). SW620 (shFOXA2) and SW620 (shControl) cells (8,000 per well) were seeded in 96-well plates. Automated phase contrast images were acquired by use of an Incucyte microscope. The Incucyte Live-Cell Imaging System provides an imbedded contrast-based confluence algorithm to compute monolayer confluence for each image and at each time point. Multiple images are collected per well and averaged to provide a representative statistical measure of confluence, thus quantifying cell growth inside the cell culture incubator.

### JQ1 experiments

For RT-qPCR or RNA-seq experiments, cells were plated at ∼0.5×10^6^ cells per well in 6-well plates and treated with 1 μM JQ1 or 0.1% DMSO (v/v) control for 24 h, then cells were lysed and RNA was extracted with TRIzol (Ambion Cat. No. 15596018) according to the manufacturer’s protocol. RNA quality was measured by using the Agilent 2100 Bioanalyzer (Agilent Technologies, USA). RNA with RIN (RNA integrity number) >7 was used for RT-qPCR or RNA sequencing.

### Tissues microarray and immunohistochemistry (IHC)

The human CRC tissue microarray (Shanghai Outdo Biotech Cat. No. HLin-Ade075Met-01) containing 75 cases of normal colon tissues, primary CRC tumors and metastases was purchased for IHC to determine FOXA2 expression. Of the 75 cases, there were 27 primary tumors and 18 liver metastases. Matched pairs of primary tumors and liver metastases are 18. IHC analysis was performed by Outdo Biotech (Shanghai, China) using standard techniques as described^29^. Briefly, all specimens on the tissue microarray were evaluated by H&E (hematoxylin-eosin) staining to ensure the pathological types before IHC staining. The tissue microarray was probed using the primary antibody (1:4,000 dilution) against FOXA2 (Abcam Cat. No. ab108422). The degree of immunostaining was scored by two independent investigators without prior knowledge of the clinical data. The IHC scores were calculated as previously described^29^. In brief, more than 1,000 tumor cells for each sample were analyzed under microscope. The percentage of positively nuclear stained tumor cells was recorded and varied from 0%-100%. FOXA2 expression was quantified using a grading system based on the percentage of FOXA2-positive cells, and the scores were as follows: 0, < 5%; 1, 5%-24%; 2, 25%-49%; 3, 50%-75%; 4, > 75%. The intensity of nuclear staining was scored as follows: 0, no staining; 1, weak staining; 2, moderate staining; 3, strong staining. A final IHC score was calculated as the product of the score of “percentage of FOXA2-positive cells” and “intensity of staining”. All primary data are shown in Supplementary Table 3.

### Mouse model

Mice were housed in facilities managed by the Tsinghua University Animal Resources Center and all experiments were performed in accordance with Tsinghua University’s Animal Care and Use Committee guidelines. SW620 cells were under domestication in liver for three times. Briefly, we used 6-week-old female Balb/c nude mice to domesticate SW620 cells. About 3 weeks later after intrahepatic injection, the mice burdened with liver tumors were sacrificed. Liver tumors were then resected and minced under sterile conditions. The minced tissues were placed in DMEM medium with 100U/ml collagenase and hyaluronidase for 1.5 hours at 37°C and then filtered by 200-mesh filter followed by centrifugation. Next, cells were resuspended and grown in DMEM medium with 10% FBS. Intrahepatic injection was performed as described^26^. Briefly, female Balb/c nude mice, 5-8 weeks, were used for surgery. Mice were anesthetized with avertin (From Tsinghua University Animal Resources Center). The skin was incised and tumor cells (5×10^6^) with shFOXA2 or shControl in 40 μl PBS were injected into the right liver lobe under the capsule. Mice were killed ∼6 weeks later, and the number of liver metastases and the metastatic area were quantified. Subcutaneous injection of tumor cells was performed as described^26^. Briefly, tumor cells (2×10^6^) were resuspended in 40 μl PBS and were injected subcutaneously into the right flank of nude mice for 1 injection site per mouse. Six days after injection and every 2 days thereafter, the length and width of tumors were measured. Volume was calculated as length*width^2^/2^30^.

### RNA-seq library preparation and sequencing

RNA-sequencing (RNA-seq) libraries were prepared by using the NEBNext Ultra Directional RNA Library Prep Kit for Illumina (NEB Cat. No. E7420), according to the manufacturer’s instructions. The sequencing was performed by Hiseq1500 (Illumina).

### RNA-seq and microarray data analysis

The fastq files from RNA-seq experiments were mapped to the human genome (hg19) by using STAR^31^ with parameters –outFilter-ismatchNoverLmax 0.05. To measure expression, we calculated the raw counts for each gene by using the analyzeRepeats command from HOMER (http://homer.salk.edu/homer/)^32^ with the option “rna” and the default parameters. We identified genes with differential expression between SW480 and SW620 cells by using edgeR^33^ with several criteria (|log2fc| ≥1, logCPM ≥1 and false discovery rate [FDR] ≤ 0.05).

For published microarray (GSE41258^34^, GSE49355^35^, GSE6919^36^, GSE32269^37^, GSE85258^38^ and GSE59000^39^) and RNA-seq dataset (GSE50760^40^), we downloaded the normalized expression values directly from Gene Expression Omnibus (GEO) database.

### ChIP-seq library preparation and sequencing

About 5-10×10^6^ crosslinked cells were used for ChIP-seq, as described^41^. After crosslinking, chromatin was fragmented by sonication, and the mixture was purified with magnetic beads (Millipore Cat. No. 16-157) conjugated to 1ng of the antibodies against H3K4me1 (Abcam Cat. No. ab8895), H3K4me2 (Millipore Cat. No. 07-030), H3K4me3 (Millipore Cat. No. 07-473), H3K27ac (Abcam Cat. No. ab4729) or FOXA2 (Proteintech Cat. No. 22474-1-AP). ChIP-sequencing libraries were prepared by using the NEBNext Ultra DNA Library Prep Kit for Illumina (NEB Cat. No. E7370), according to the manufacturer’s instructions. After barcoding, pooled DNA was sequenced (HiSeq 1500, Illumina) to achieve a minimum of 1×10^7^ aligned reads per sample.

### ChIP-seq data analysis

Fastq files from ChIP-seq experiments were mapped to the human genomes (hg19) by using STAR^31^ with parameters –outFilter-ismatchNoverLmax 0.05. For the ChIP-seq of histone modification, enriched loci were identified by using the findPeaks command from HOMER (http://homer.salk.edu/homer/)^32^ with the option –style histone, 4-fold enrichment over the input sample, 4-fold enrichment over local background, at FDR = 0.001, and normalization to 10 million mapped reads per experiment.

Peaks of H3K4me2 upstream and downstream 3 kb away of gene’s TSS were defined as enhancers. Super-enhancers were identified using the original strategy used by the Young lab^20^ using our H3K27ac data. First, peaks are found just like any other ChIP-Seq data set. The peaks found in ChIP-seq of histone modification within a given distance are ‘stitched’ together into larger regions (by default over 12.5 kb). The super-enhancer signal of each of these regions is then determined by the total normalized number reads minus the number of normalized reads in the input. These regions are then sorted by their score, normalized to the highest score and the number of putative enhancer regions, and then super-enhancers are identified as regions past the point where the slope is greater than 1.

Genomic binding peaks for transcription factor FOXA2 were identified with option -style factor, 2-fold enrichment over the input sample, 2-fold enrichment over local background, at the false discovery rate of 0.001.

All the peaks of histone and transcription factor were annotated to the nearest TSS (transcription start site), TTS (transcription termination site), Exon (Coding), 5’ UTR Exon, 3’ UTR Exon, Intronic, or Intergenic of genes using annotatePeaks.pl command.

All peaks between the different cell types per comparison were merged into one peak set by using mergePeaks –size given. To obtain differentially bound peaks, tags were counted from each experiment by using getDifferentialPeaks and were considered significant with default parameters (4-fold difference and *P* value = 0.0001). These enhancer regions with peak signals >4-fold in SW620 than in SW480 were named “SW620-unique” regions; the opposite regions were named “SW480-unique” regions. The similarly bound peaks were determined by using the same option. All peaks which were not part of either the differential bound or similar bound peaks were filtered out.

### *Do novo* motif finding and known motif enrichment

We used HOMER^32^ with *de novo* motif discovery and known motif enrichment. Motif finding for histone modification (H3K4me2, H3K27ac) regions distant from 3 kb up-and down-stream of TSS was performed on sequence from 1000 bp relative to the peak center, whereas motif finding for transcript factors (FOXA2) was performed on sequence of given length. Briefly, sequences were divided into target and background sets. Background sequences were then selectively weighted to equalize the distributions of G/C content in target and background sequences to avoid comparing sequences of different general sequence content. Motifs of 8, 10 and 12 bp were identified separately by first exhaustively screening all oligonucleotides for enrichment in the target set compared with the background set by using the cumulative hypergeometric distribution to score enrichment. Up to two mismatches were allowed in each oligonucleotide sequence to increase the sensitivity of the method. Top oligonucleotides for each length with the lowest *P* values were then converted into probability matrices and heuristically optimized to maximize hypergeometric enrichment of each motif in the given data set. As optimized motifs were found, they were removed from the dataset to facilitate the identification of additional motifs in subsequent rounds. HOMER also screens the enrichment of known motifs previously identified by analysis of published ChIP-ChIP and ChIP-Seq data sets by calculating the known motifs’ hypergeometric enrichment in the same set of G/C normalized sequences used for de novo analysis. Sequence logos were generated using WebLOGO (http://weblogo.berkeley.edu).

### Gene Set Enrichment Analysis

We defined tissue-specific gene sets by using the expression values yield by the Genotype-Tissue Expression (GTEx) project. We downloaded the file that contained the median RPKM of each gene by tissues and cell types from the latest release version, V6p. For each gene, we ranked the median expression value for each tissue or cell type in decreasing order. Genes defined as tissue-specific needed to meet two criteria: 1) ranked in the top 5 among all tissue and cell types and 2) also highly expressed (∼90th percentile of all genes) in particular tissues and cell types. The log2 ratios were computed for several datasets from normalized expression data, including the primary versus liver metastatic tumor samples from GSE41258^34^, GSE49355^35^, GSE50760^40^, GSE6919^36^, GSE32269^37^, GSE85258^38^ and GSE59000^39^ datasets, and the DMSO versus JQ1 treatment on SW620 cells. Then, the pre-ranked Gene Set Enrichment Analysis^17^ was performed with tissue-associated gene sets and dataset with default parameters. We show the scatter plot of FDR (q-values) versus normalized enrichment score (NES) for each analysis.

### Removing expression noises caused by contamination of host organ

In order to further confirm the tissue-specific gene signatures we found in the patient samples, we removed differentially expressed probes/genes that could be introduced by the contamination of distant organ (i.e. normal liver tissue) in the metastasis samples. We adapted a published method^35^ to remove ambiguous probes/genes. Because hepatic metastasis (HM) sample may be contaminated by certain normal hepatic tissue (HN), the measured expression value of HM for each probe/gene, *mHM*, is different from the real expression value:

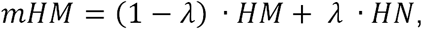

where *λ* is the ratio of HN contamination in HM. Thus, we removed the ambiguous up-regulated probes/genes between hepatic metastasis sample (HM) and primary colon tumor (CT) using two criteria:

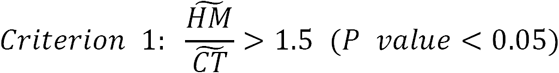

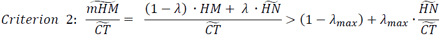

where 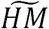 and 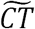 are the median expression values of multiple samples in the same data set. We assume the maximum contamination ratio is 20% (*λ*_*max*_ = 0.2). The results of different values of *λ*_*max*_ are shown in Extended Data Figure 8 as well. For datasets (GSE49355, GSE50760) without normal liver tissue (HN), the expression values of probes/genes in HN were calculated using corresponding samples in GSE41258 (array) or GTEx database (RNA-seq).

We examined the enrichment of tissue-specific gene signatures with odds ratios calculated from four numbers (Figure 1f, Extended Data Figure 8): (1) total number of genes; (2) number of tissue-specific gene signatures (i.e. liver specific genes defined by GTEx database); (3) number of up-regulated genes filtered by the above two criteria; (4) number of liver specific genes filtered by the above two criteria. *P* value of each odds ratio is calculated using Fisher’s exact test.

We used the same strategy to examine the colon-sigmoid specific gene signatures down-regulated in the metastasis samples (Extended Data Fig. 1).

### Kaplan-Meier survival analysis

We downloaded RNA-seq data for COAD patients with clinical data in the TCGA from the NCI Cancer Genomics Hub (CGHub)^42^. For overall survival (OS) analysis, the event call was derived from the ″*vital status*″ parameter. The *time_to_event* is in days, equal to *days_to_death* if the patient died; if the patient is still living, the time variable is the maximum (*days_to_last_known_alive, days_to_last_followup*). For recurrence-free survival (RFS), the event call was derived from the ″*new_tumor_event_after_initial_treatment*″ parameter. The *time_to_event* is in days, equal to maximum (*days_to_new_tumor_event_after_initial_treatment, days_to_tumor_recurrence*) if there is an event. In the Kaplan-Meier survival analysis, 433 and 203 patients were used for OS and RFS analysis, respectively. We first divided patients into two groups (bottom 20% and top 20%) by median expression level of signature genes, then used Kaplan-Meier survival analysis^43^ to analyze OS and RFS analysis via the survival package (https://cran.r-project.org/web/packages/survival) in the R environment for statistical computing and computed significance with the log-rank test.

